# Y665F variant of mouse *Stat5b* protects against acute kidney injury through transcriptomic shifts in renal gene expression

**DOI:** 10.1101/2025.02.19.639141

**Authors:** Jakub Jankowski, Hye Kyung Lee, Lothar Hennighausen

## Abstract

The impact of single nucleotide polymorphisms (SNP) on physiology is often underestimated. One amino acid change can result in a variety of phenotypes apparent only in response to disease or injury. Even known pathogenic SNPs have widespread effects that are currently unaccounted for. In this study, we investigated the impact of the known activating and pathogenic *STAT5B^Y665F^* mutation in a renal injury context in mice carrying this variant. Using ischemia-reperfusion model of acute kidney injury, immunohistochemistry, RNA-seq and ChIP-seq, we establish the protective role of STAT5B activation in renal epithelium and showcase the shifts in transcriptomic landscape in a tissue not associated with the usual human phenotype of the *Stat5b^Y665F^* mutation. Our data indicate new links between the JAK/STAT pathway and known kidney injury markers, contribute to the understanding of the sexual dimorphism of renal disease, and provide new potential targets for JAK inhibitor– and amino acid transport modulation-based therapies.

## INTRODUCTION

Disease-associated single nucleotide polymorphisms (SNPs) are usually viewed through the lens of their most prominent phenotype. However, focusing an investigation on only one disease, organ, or tissue, results in overlooking their effect on the whole system and their contribution to a range of physiological responses. Further, transcriptional patterns shift significantly in any pathological setting, often putting additional strain on the altered protein’s activity. Thus, limiting inquiry to the most common symptoms is more often than not short-sighted.

Single missense mutation can significantly alter protein’s structure, and, albeit with exceptions, most proteins are not restricted to one tissue.^1–3^ Transcription factors, proteins directly interacting with DNA and regulating gene expression, are usually ubiquitous. Any introduced amino acid change can result in widespread consequences, even if they are semi-tissue-restricted. For example, the same intronic SNP of the HNF1b transcription factor, mostly expressed in kidney and pancreas, can enhance susceptibility to type-2 diabetes development, but also increase risk of prostate cancer, and missense mutations in TBX5, usually associated with cardiomyocytes, can result in Holt-Oram syndrome and abnormal upper limb development.^4–6^

STAT transcription factors are not an exception. They perform diverse functions, both in homeostasis and disease, usually as a result of their phosphorylation due to extracellular stimuli, like cytokines. Point mutations in STATs are linked to a variety of phenotypes, spanning all the major organ systems.^7–11^ Several mouse models harboring STAT mutations have been developed, establishing their role as crucial for body function. For example, STAT1 knockouts are extremely susceptible to infection and die within days after birth if not in housed in pathogen-free conditions, while STAT3 knockouts are not viable.^12^ ^13^ Multiple STAT knockout mouse strains have been investigated, but very few of them try to recapitulate gene variants observed in human disease. Among them, mut-Stat3 strain mimics human hyper-IgE syndrome and STAT3 G421R strain displays T-cell dysregulation aiming to mimic primary immune deficiencies.^14,15^ We have recently developed a *Stat5b* mutant mouse line, in which we introduced a SNP changing tyrosine to phenylalanine in position 665 (Y665F).^16^ This mutation has been detected in multiple leukemia patients and was hypothesized to be pathogenic due to STAT5B hyperactivation.^17,18^ The mice, while displaying number of hematopoietic abnormalities, did not directly develop malignancies. They did, however, show severe differences in the development of mammary glands, stemming from altered enhancer activity and downstream gene expression, among other physical abnormalities.^16^ There are only a few pieces of evidence for *Stat5b* being a key injury response regulator. Because of the heterogeneity of both the cellular components of the kidney and renal injury etiologies, its role is inconclusive. While impaired JAK/STAT5 signaling contributes to the development of polycystic kidney disease and chronic kidney disease, podocyte STAT5 ameliorates focal segmental glomerulosclerosis.^19–21^ Only one poster publication indicates protective role of STAT5 in acute, cisplatin-induced injury model.^22^

In this manuscript, we follow up on our previous studies to investigate the extent of the transcriptomic shifts caused by the *Stat5b^Y665F^*mutation in an acute kidney injury (AKI) setting. There is no previously published data available indicating *Stat5b’s* involvement in renal ischemic injury, providing us with an opportunity to both establish its importance in AKI, and to investigate the extent of one SNP’s effect on the transcriptional landscape of renal epithelium. To accomplish that, we use a well-established mouse ischemic injury model, followed by immunohistochemistry and RNA-seq analysis, as well as investigation of our previously published ChIP-seq data. Our analysis strongly indicates that STAT5b activity regulates the severity of renal injury through multiple mechanisms, such as modulating inflammation, sex-specific gene expression and amino acid transport.

## METHODS

### Mice

#### General care

All animals were housed in the same environmentally controlled room (22–24 °C, with 50 ± 5% humidity and 12 h/12 h light–dark cycle) and handled according to the Guide for the Care and Use of Laboratory Animals (8th edition) and all animal experiments were approved by the Animal Care and Use Committee (ACUC) of National Institute of Diabetes and Digestive and Kidney Diseases (NIDDK, MD) and performed under the NIDDK animal protocol K089-LGP-23. All mice were 12-16 weeks old at the time of respective experiments. Mice harboring the *Stat5b^Y665F^* mutation (Y665F) were generated as described previously.^16^ Due to disease phenotypes requiring euthanasia at 2-3 months of age present in homozygous mice, heterozygous mutants were used.

#### Ischemia-reperfusion surgery

Randomized litters of heterozygous and wildtype-bred mice were used for all the procedures. We chose approximately 2-3 months old mice to balance avoiding mortality in our severe model (3.7% or 1/27 for data presented), while aiming for more pronounced injury than in young animals. To perform warm renal ischemia-reperfusion, in random order and with the surgeon blinded to genotype, but not sex, mice were anesthetized with ketamine/xylazine mix (100mg/kg and 10mg/kg respectively). Hair was removed from the mouse retroperitoneal area using sterilized electrical clippers and skin was cleared and prepared using betadine and ethanol swabs. Next, the mice were placed over temperature-controlled heating pad maintained at 38°C. Core temperature of the mice was sustained at approximately 35.5 – 36°C as measured by a rectal probe. Renal Ischemia was induced by clamping the renal artery for 30 minutes bilaterally. Then, the clamp was removed and skin was closed using sterile wound clips, which were removed at the time of euthanasia. Finally, the animals were injected with 1ml saline to replenish fluids, provided analgesia (sustained release Buprenorphine, 1mg/kg), and allowed to recover in a cage heated to approximately 37°C until anesthesia wore off.

### Renal Injury Assessment

Serum creatinine with a colorimetric kit (Diazyme) according to manufacturer’s instructions. For histology assessment, kidneys were fixed in 10% neutral buffered formalin for 24 hours, washed and stored in 70% ethanol. Preparation of paraffin slides and H&E staining was performed by Histoserv. Keyence BZ-9000 microscope was used to take serial, randomized photographs of renal cortex and outer medulla at total 400x magnification. At least 6 photographs were taken per animal, visualizing both kidneys to ensure uniform injury. Photographs with at least 90% tissue coverage were then overlaid with a randomized point grid and percentage of injured tubules was quantified. Approximately 350 tubules per animal were assessed. Tubular dilation, loss of nuclei or membrane integrity, and protein casts were the positive injury criteria.

### Bulk RNA Sequencing (Total RNA-seq) and Data Analysis

Bulk RNA was extracted from whole frozen renal tissue from wild-type and mutant mice and purified with RNeasy Plus Mini Kit (Qiagen, 74134). Ribosomal RNA was removed from 1 μg of total RNAs, and cDNA was synthesized using SuperScript III (Invitrogen). Libraries for sequencing were prepared according to the manufacturer’s instructions with TruSeq Stranded Total RNA Library Prep Kit with Ribo-Zero Gold (Illumina, RS-122-2301), and paired-end sequencing was done with a NovaSeq 6000 instrument (Illumina). Read quality control was done using FastQC (Babraham Bioinformatics) and Trimmomatic.^23^ RNA STAR was used to align the reads to mm10 genome. HTSeq and DeSeq2 were used to obtain gene counts and compare genotypes.^24,25^ Genes were categorized as significantly differentially expressed with an adjusted p-value below 0.05. Differentially expressed genes were visualized with ComplexHeatmap R package.^26^

### Immunohistochemistry

Kidneys were fixed in 10% neutral buffered formalin for 24 hours, washed and stored in 70% ethanol. Preparation of unstained paraffin slides was performed by Histoserv. Slides were rehydrated by washing twice in xylenes, 1:1 xylenes-ethanol solution, twice in 100% ethanol, 95% ethanol, 70% ethanol, 50% ethanol, twice in water; each washing step lasting 3 minutes. Next, slides were boiled in Antigen Unmasking Solution (Vector Laboratories) for 10 minutes and incubated in fresh 3% hydrogen peroxide for 10 minutes. After washing with water, blocking in 2.5% goat serum (Vector Laboratories) was performed for 1 hour at room temperature, after which the sections were incubated in primary antibody detecting mouse STAT5b (Invitrogen #13-5300), phospho-STAT3 (Cell Signaling #9145) diluted 1:200 in serum, or in serum alone (secondary antibody control) overnight at room temperature. Next, slides were washed twice in water and suspension of the secondary antibody was added (Goat anti-mouse IgG, Vector Laboratories, MP-7452 or goat anti-rabbit IgG, Vector Laboratories, MP-7451). After an hour of incubation at room temperature, slides were washed, and DAB substrate (Vector Laboratories) was added for 10 minutes. After thorough washing with water, hematoxylin solution (Sigma-Aldrich) was added for 3 minutes, followed by another wash. Slides were dehydrated by following the reverse order of initial washes, ending with xylenes and subsequent mounting with Permount (Fischer Chemical). Additionally, CD3 staining was performed by Histoserv Inc., using anti-mouse CD3 antibody (Cell Signaling Technology #78588). To quantify CD3-positive Keyence BZ-9000 microscope was used to take serial photographs of renal tissue at total 200x magnification. At least 4 photographs with at least 90% tissue coverage were assessed for each animal, and number of CD3-positive cells per microscope field was averaged per animal.

### Statistics and Reproducibility

GraphPad PRISM 10 was used to analyze experimental data. Normal distribution test (Shapiro-Wilk) was performed before assessing statistical significance of the findings by using appropriate measure detailed in each figure description. Tukey’s method was used for multiple comparison correction. All tests used two-tailed p-value and statistical significance was set at p<0.05. Levels of statistical significance were described on graphs as follows: * *P* < 0.05, ** *P* < 0.01, *** *P* < 0.001, **** *P* < 0.0001. Group sizes are reported in detail in figure descriptions. Error bars on graphs represent standard error of the mean (SEM). Any statistically analyzed data is derived from biological replicates.

## RESULTS AND DISCUSSION

### *Stat5b^Y665F^* mutation protects male mice against renal injury

To establish the effects of the *Stat5b*^Y665F^ mutation on renal injury, we used a 30-minute bilateral ischemia-reperfusion model and euthanized mice at the 24-hour timepoint. Our primary readout of the injury efficacy and response was plasma creatinine increase (Fig. 1a). We observed smaller rise in creatinine in heterozygous mutant males compared to WT (mean 0.4948 vs. 1.231, p<0.01), but not in females (mean 0.7867 vs. 1.038, p>0.05). As female mice are often regarded as more protected from acute renal injury, we purposefully used a severe model to elicit a measurable response. Following, we investigated renal histology of male mice and confirmed the protective effect of the Y665F mutation, as fewer tubules were visibly injured in mutants than WT mice (Fig. 1b, c).

**Fig. 1.**
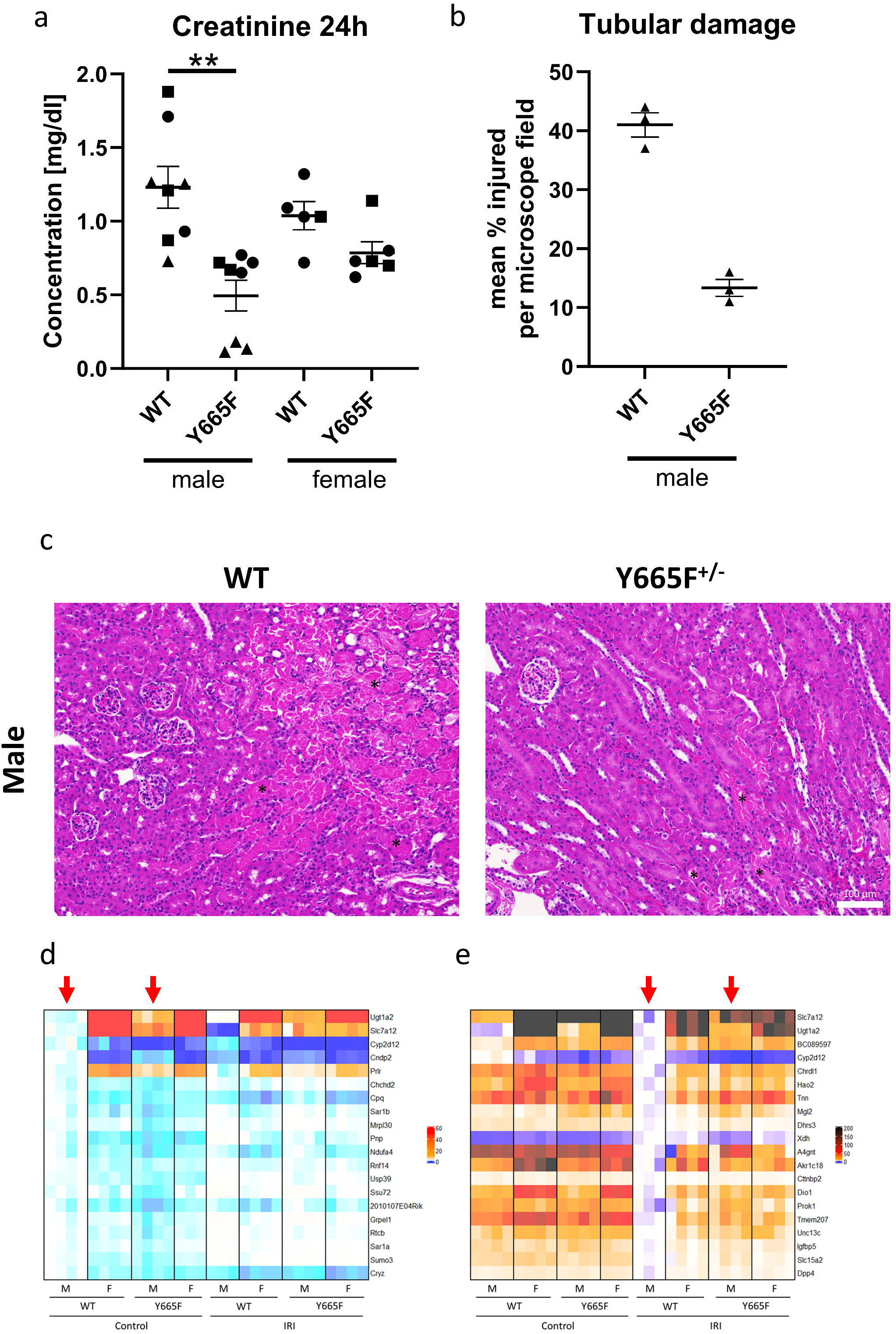
Male *Stat5b^Y665F^*mutants are protected from ischemia-reperfusion injury. Serum creatinine (a) and tubular injury quantification (b) at 24 hours after 30’ bilateral ischemia-reperfusion surgery in male and female wild type and Y665F mice. Representative images of renal tissue from male mice 24 hours after injury, with example injured tubules marked with asterisks (c, d). Heatmaps of 20 most significantly deregulated genes between male wild type and Y665F mice at the baseline (e) and after injury (f). a – n = 8 for male and n = 6 for female mice (one WT female did not survive to the 24h timepoint and was omitted), Two-way ANOVA with group mean comparisons, Bar = SEM, ** *P* < 0.01, different symbols denote surgery cohorts;, b – n = 3, average of injured tubules counted in randomized photographs; c – n = 3, bar = 100 μm, 400x magnification; d, e – red arrows indicate columns compared for statistically significantly different gene expression.

Gene expression of *Stat5b*, measured by bulk RNA-seq, did not change in the Y665F strain, not only at the baseline, which is in line with our previous observations^16^, but also in the injury setting (Supplementary Fig.1a). Antibody staining revealed STAT5b-positive nuclei, but no evident differences between experimental groups (Supplementary Fig. 1b). This suggests that the germline *Stat5b*^Y665F^ mutation could be preconditioning kidneys to resist injury, rather than activating STAT5b to act acutely during ischemia and reperfusion. Expanding the investigation to the entire STAT family, we identified *Stat3* as the only other gene displaying increase in expression after injury, smaller in mutant males compared to WT (Supplementary Fig. 2a, Supplementary Spreadsheet 1). While we saw an increase in phosphorylated STAT3 after injury, once again we did not observe a clear quantitative difference between WT and mutant mice (Supplementary Fig. 2b).

Next, we investigated whether infiltrating T-cells could have a significant impact on the injury outcomes in our model. Y665F mutants, both male and female, present with splenomegaly (Supplementary Fig. 3a). CD3 antibody staining revealed significantly more T-cells present in the renal tissue before injury, but uniformly decreasing after injury (Supplementary Fig. 3b,c). This suggests that while the increased baseline can be to a certain degree protective in the injury setting, no abundant additional T-cell infiltration occurred at the 24h timepoint.

In consequence, we hypothesize that epithelial STAT5b activity is the source of renal protection. We performed bulk RNA-seq to gain an overview of the renal transcriptional landscape before and after injury. The analysis revealed 187 significantly upregulated and 396 downregulated genes in between male mutant males compared to WT males at the baseline, while 1124 were upregulated and 678 downregulated after injury. In comparison, female mice displayed no differences in gene expression between the mutants and WT mice before injury, and only six genes reached statistical significance after ischemia-reperfusion (Fig. 1d, e, Supplementary Spreadsheet 1), suggesting significant sex-dependent effect of our observations.

### Known injury response pathways are altered by *Stat5b^Y665F^* mutation

Comparing WT and mutant males after injury, there were several deregulated genes closely linked to AKI response. *Lcn2*, one of the most commonly used kidney injury markers, is significantly more upregulated in male WT than in mutant mice after ischemia-reperfusion (Fig. 2a). *Lcn2* has been reported to interact with JAK/STAT pathway through multiple mechanisms, and is usually described as anti-inflammatory in the renal injury context. It can activate NFκB/STAT3 pathway in macrophages^27^, alleviate infection through STAT1 and STAT3 downregulation^28^, but also is itself subject to upregulation through Il-1/STAT3 mechanism.^29^ Its direct link to STAT5b is unknown and using previously published ChIP-seq data we were unable to find any *Stat5b* binding peaks lining up with GAS motifs (characteristic for STAT proteins) or H3K27ac marks designating active chromatin (Supplementary Fig. 4a).

**Fig. 2.**
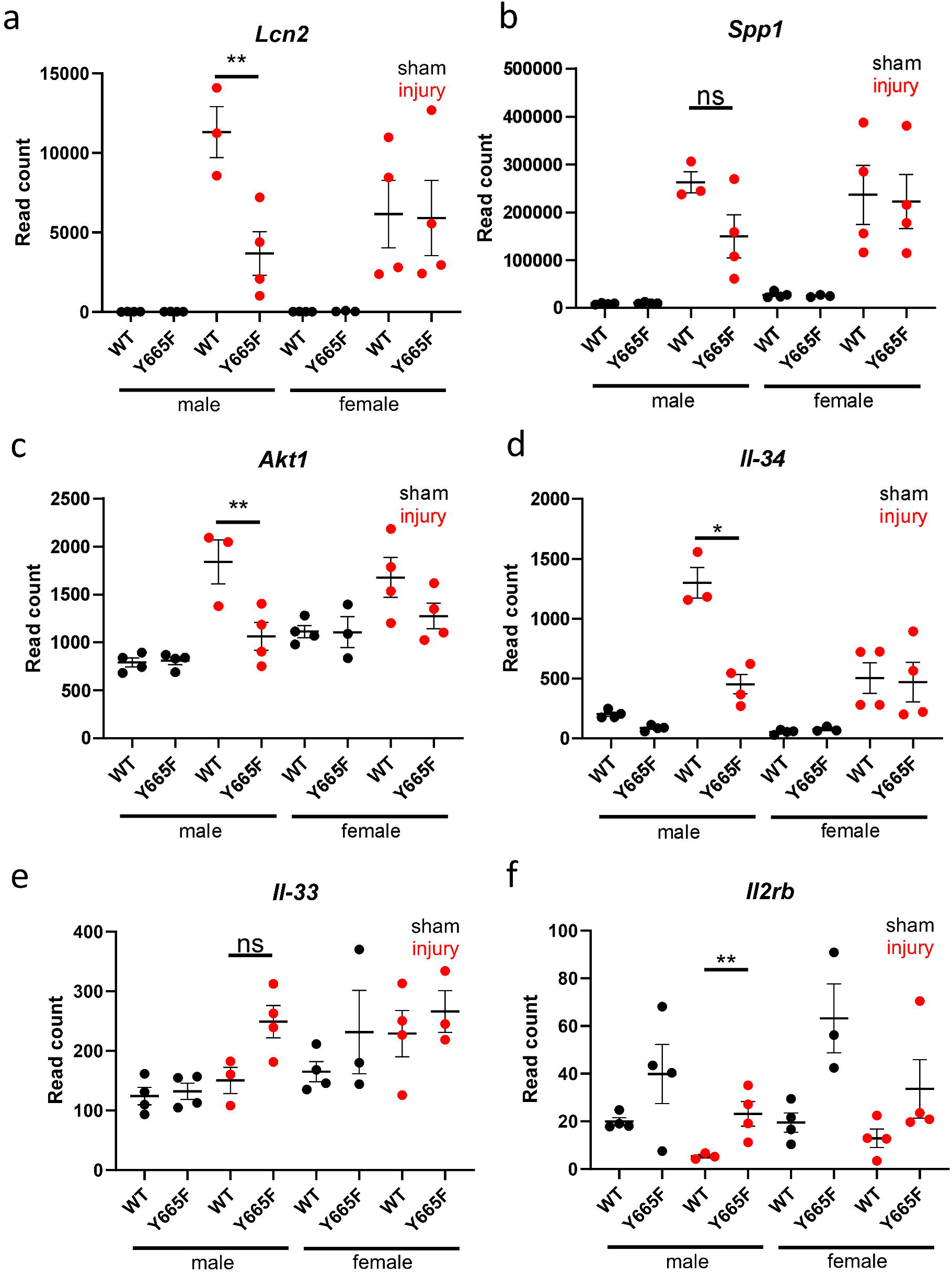
Expression of multiple injury-associated genes suggests attenuated renal injury in Y665F mutants. Normalized DeSeq2 reads of *Lcn2* (a), *Spp1* (b), *Akt1* (c), *Il-34* (d), *Il-33* (e) and *Il2rb* (f) in male and female wild type and Y665F mutants before and after injury. n = 3-4, Two-way ANOVA with group mean comparisons, Bar = SEM, * *P* < 0.05, ** *P* < 0.01

Ischemic injury is known to activate osteopontin (*SPP1*) expression and release.^30–32^ Although its expression is not statistically significantly decreased compared to WT in the injury setting (Fig. 2b), it possesses an upstream GAS motif occupied by STAT5b as evidenced by ChIP-seq analysis (Supplementary Fig. 4b). RNA-seq points to activation of downstream targets of *Spp1*, such as *Akt1* and *PIK3ca* (Fig. 2c).^33,34^ Using ChIP-seq, we observed several occupied GAS motifs in the intronic regions of *Akt1*, and smaller peaks at the promoter of the *PIK3ca*, which might be indirect binding sites, as no GAS motifs are present in the locus (Supplementary Fig. 4c, d). PI3K/AKT pathway is well-known to modulate renal apoptosis, autophagy and aging, but its link to STAT5b has not yet been established.^35–37^

JAK/STAT pathway is closely linked to cytokine stimulation and regulation of immune response. STATs are known regulators of interleukin production, while at the same time being under their transcriptomic control. For example, STAT1 regulates production of Il-1β during infection, but can also be induced by Il- 2 and Il-6^38,39^, and Il-6/STAT3 axis can be targeted in multiple disease settings.^40,41^ Our data indicates that STAT5b can regulate *Il-34* expression, elevated significantly higher in WT males than the mutants (Fig. 2d). Several GAS motifs and H3K27ac marks within *Il-34* locus support this conclusion (Supplementary Fig. 4e). While no strong link has been established between Il-34 and JAK/STAT signaling, there is abundant evidence suggesting that the interleukin is deleterious in ischemic setting.^42–44^ Surprisingly, we did not observe the effect of the injury on the Il-33 and Il-2, known modulators of hypoxia and renal injury (Fig. 2e, f).^45–47^ While the increase in *Il2rb* in mutant males was significantly higher than that in WT males, relatively low read count, combined with lack of occupied GAS motifs observed within the ChIP-seq data, suggests that in our model this pathway does not play a significant role. However, this observation needs to take into consideration earlier reports of T-cell STAT5b activating T-cell *Il2ra* expression.^48^ While more work is necessary, this could potentially reinforce the hypothesis that epithelium is the main source of the protective effect. Both Il-2 and Il-33 are involved in T-cell activation, and together with lack of *Stat1* gene induction, the lack of increase in expression might suggest they remain largely inert.

### Renal STAT5b drives sexually dimorphic gene expression

Response to renal disease and its progression is highly dependent on the sex of the patient).^49^ It is generally recognized that men are at higher risk of AKI, while chronic kidney disease is more prevalent in women, though its progression is slower than in men.^50,51^ The female protection against renal injury was proposed to diminish after menopause, strongly indicating that sex hormones play a role in the process.^52^ RNA-seq results displaying the most deregulated genes between males before and after injury showcase a trend towards feminization of gene renal gene expression in the Y665F mutants (Fig. 1d, e). For example, *Ugt1a2* and *Slc7a12* are expressed at higher levels in male mutants, similarly to the levels found in female mice, while expression of *Cyp2d12* and *Cndp2* decreased in mutant compared to WT males. After injury this trend becomes even more obvious, as genes with elevated expression in female mice and mutant males fail to follow in male WT mice. This observation marks STAT5b activity as driver of sexually dimorphic gene expression. A similar effect has been reported in the liver and skeletal tissue, indicating them as potential targets of follow-up studies and tissues of concern in patients harboring the *STAT5B^Y665F^*mutation.^53–56^

To try to discern why the sexually dimorphic effect is present, we investigated known hormone regulators present in the kidney. Estrogen receptors and sirtuins, acting in tandem, are known protective factors in renal injury and chronic disease setting.^57,58^ We observed elevated *Esr1* transcript in Y665F males, but not the females, both before and after injury (Fig. 3a). Estrogen receptors can activate STAT5b through phosphorylation by JAK2^59,60^, but also a region in a small radius around *Esr1* presents several occupied GAS motifs, suggesting a feedback loop (Supplementary Fig. 5a). ESRα-STAT5b axis plays a role in mineral and water homeostasis and renal cell carcinoma development, but its role in renal injury is to be determined.^61,62^ In comparison, there is little recent information about the role of prolactin in kidney disease. Prolactin receptor (*Prlr*) expression was one of the most significantly deregulated genes between male experimental groups, and elevated in Y665F mutants (Fig. 3b). Interaction between prolactin and its downstream effectors like cyclin D1 and TP53 (Fig. 3c, d) is usually explored in the context of mammary tumors. Our data might indicate that in the renal injury setting, prolactin promotes cellular proliferation and regeneration.^63–65^

**Fig. 3.**
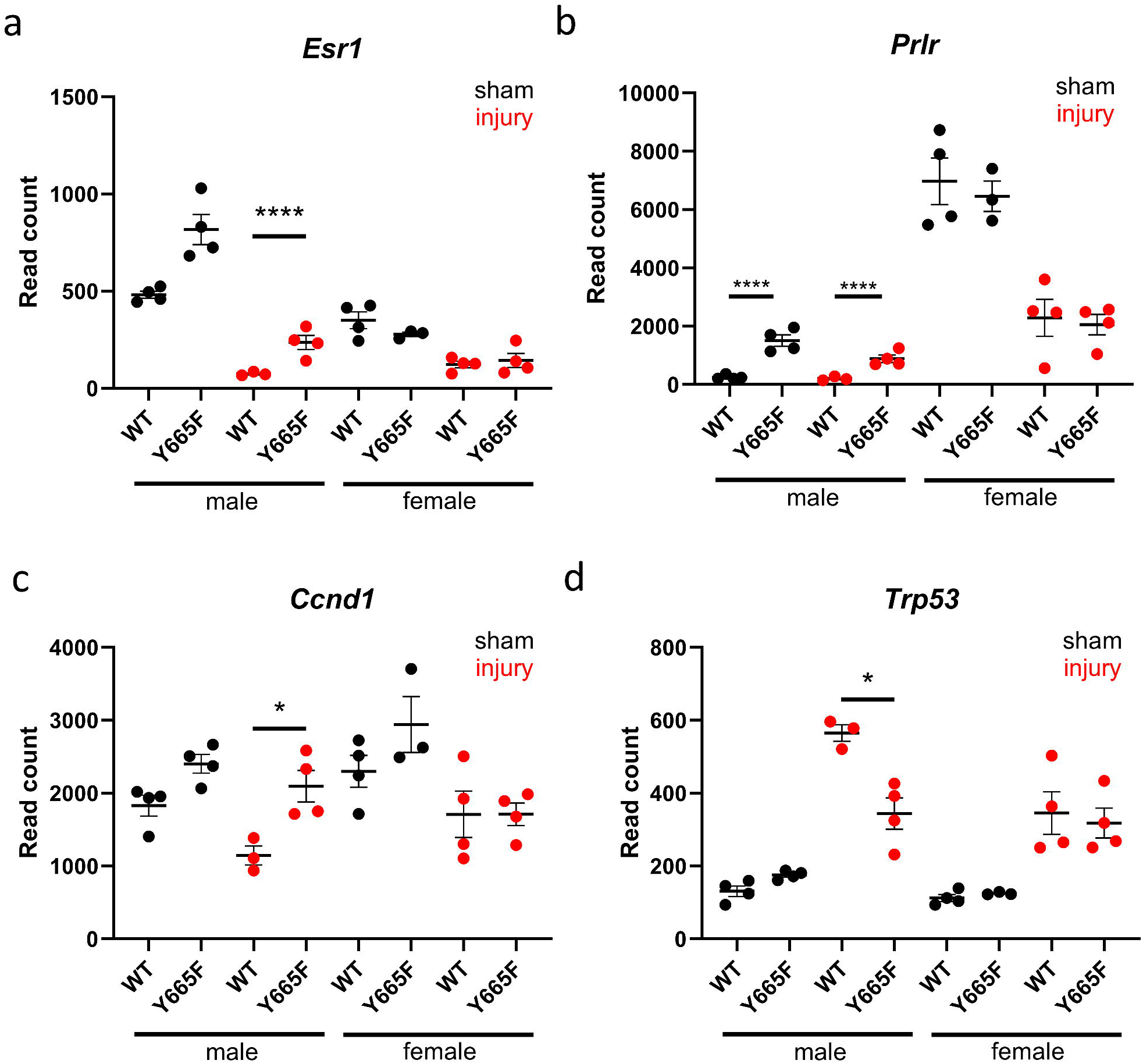
Sex hormone activity is dysregulated in Y665F males and promotes injury resistance. Normalized DeSeq2 reads of *Esr1* (a), *Prlr* (b), *CCND1* (c) and *Trp53* (d) in male and female wild type and Y665F mutants before and after injury. n = 3-4, Two-way ANOVA with group mean comparisons, Bar = SEM, * *P* < 0.05, **** *P* < 0.0001

Another result of the *Stat5b^Y665F^* mutation worth considering is the apparent feminization of gene expression on chromosome X (Fig. 4). There is a number of X-linked renal diseases, clinically more prevalent in men than women, and aberrant gene expression resulting from disruption of the JAK/STAT pathway is worth investigating.^66^ Among 59 differentially expressed genes in male injury group, 12 possess STAT5b peaks occupying GAS motifs and thus are directly activated or inhibited by it (Supplementary Fig. 5b, c). Notably, majority of those genes does not display sexual dimorphism prior to the injury, explaining which could be the subject of a follow-up study.

**Fig. 4.**
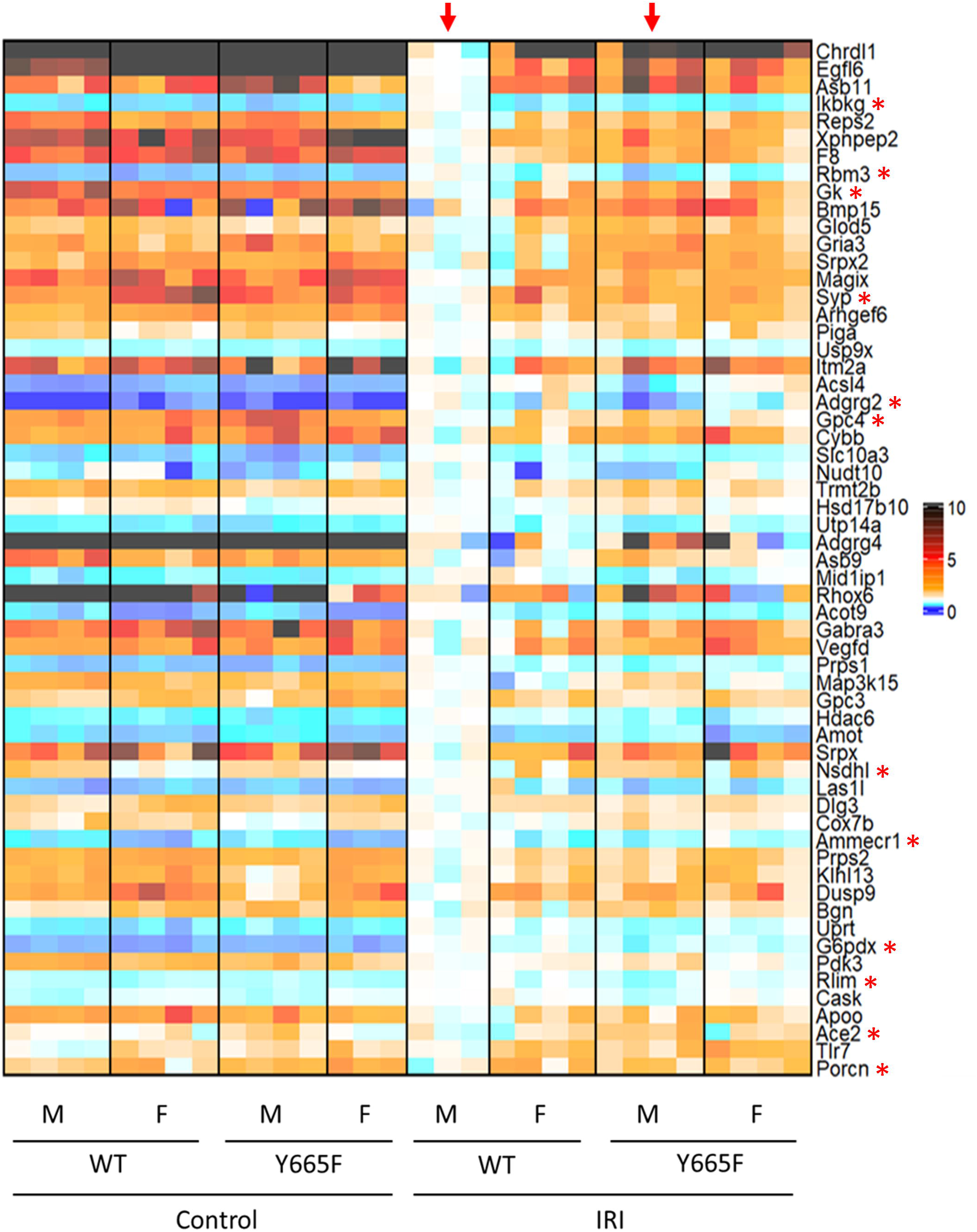
Injury causes differential expression of X-chromosome genes in wild type and mutant males. Heatmap of differentially expressed genes comparing wild type and mutant male mice after injury (arrows). Genes present on chromosome X are visualized. Asterisks indicate genes with STAT5b peaks present in ChIP-seq analysis suggesting direct modulation.

### STAT5b drives renal amino acid transport

Amino acid transport and resorption is a key function of the kidney.^67^ Renal epithelium expresses a vast number of transporter proteins, most notably of the SLC family. As such, modulating and preserving those critical mechanisms has been a target of several therapeutic approaches, like inhibition of the SLC6A19 protein to attenuate renal injury.^68,69^ ACE2 inhibition, often used in clinical setting, also can affect epithelial amino acid transport. Numerous SLC family members, *Ace2*, and several others were found significantly deregulated in male WT mice compared to mutants after injury (Fig. 5). While the JAK/STAT pathway is linked to amino acid transport, direct link to STAT5b signaling has not been established thus far.^70–72^

**Fig. 5.**
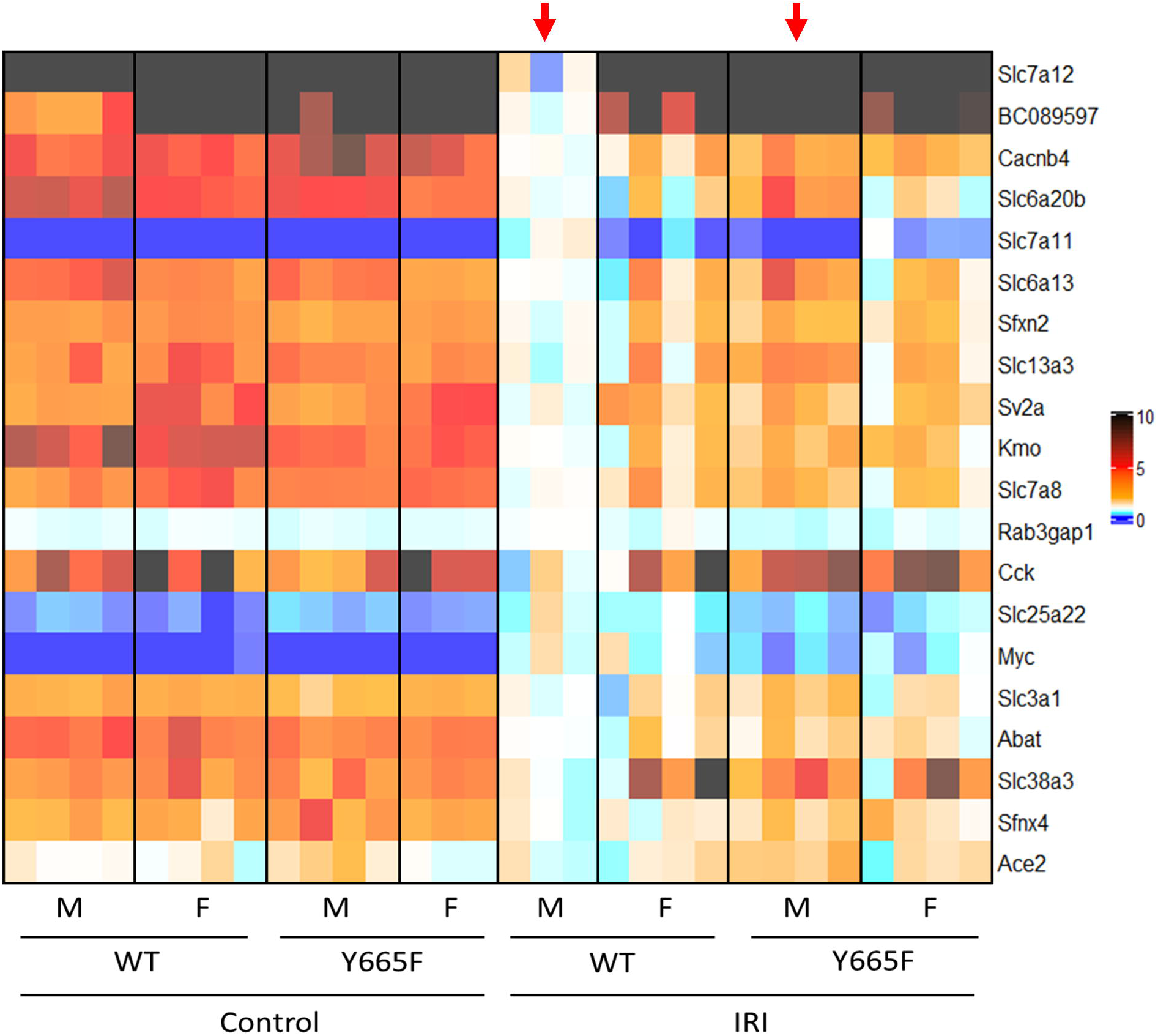
Amino acid transport in Y665F mice is altered after injury. Heatmap of differentially expressed genes comparing wild type and mutant male mice after injury. Top 20 genes categorized as taking part in amino acid transport by the GSEA Molecular Signal Database are presented.

### Summary

In this manuscript, we have established the role of the *Stat5b^Y665F^*variant as modulator of renal injury. Attenuated elevation of plasma creatinine in compared to controls, histology, and injury marker expression confirms that STAT5b activation helps protect against kidney ischemia-reperfusion model. The lack of injury-stimulated changes in *Stat5b* mRNA level and abundance of STAT5B nuclear antibody staining, as well as increased immune infiltration at the baseline, suggests prolonged presence of adaptive mechanisms allowing for the observed protection. We present several possible mechanisms explaining the findings. First, the expression levels of a number of inflammatory marker genes, including *Lcn2*, *Il-34* and *Spp1,* were lower in mutants compared to those in WT mice. Second, there is a significant sexually dimorphic component in any renal injury model, here enhanced due to the effect of hyperactivated STAT5b skewing male renal gene expression pattern towards female. Another aspect of this change is differential expression of X-chromosome genes, including ones directly under *Stat5b* control. Lastly, we show altered amino acid transporter expression, which is a potential target of already existing therapies, including those modulating ACE2 activity. Finally, our work for the first time discusses the role of STAT5b in acute ischemic injury. While JAK inhibition is used to alleviate symptoms of chronic and diabetic kidney injury, modulating STAT5b activity may be beneficial to treat immediate injury response.^73–75^

## DATA AVAILABILITY STATEMENT

All RNA-seq datasets generated for this study were deposited in Gene Expression Omnibus (GEO) with the accession number GSE289432 (Reviewer token: otkbiusujzibtil). In addition to original data, GEO series GSE114292 and GSE270652 (Reviewer token: epejwmwgblclbof) was used to investigate ChIP-seq loci of STAT5b-regulated genes. Further, data necessary to replicate figures, including original photographs, was deposited in Zenodo data sharing repository with the DOI: 10.5281/zenodo.14862695. Any additional data or materials are available on request.

## ACKNOWLEDGEMENTS

We thank the NHLBI Genomics Core for performing NGS, Jeff Reece and the Advanced Light Microscopy & Image Analysis Core for microscopy advice and equipment, and Chengyu Liu and NHLBI Transgenic Core for generating the mice. This work utilized the computational resources of the NIH HPC Biowulf cluster (http://hpc.nih.gov).

## AUTHOR CONTRIBUTIONS

Conceptualization and methodology: J.J., H.K.L., and L.H.; Formal analysis and validation, data curation, and visualization: J.J; Investigation: J.J.; Resources: L.H.; Writing – original draft: J.J.; Writing – review and editing: J.J., H.K.L., and L.H.; Supervision, administration, and funding acquisition: L.H. All authors approved the final version of the manuscript.

