## Supplementary Figures for "Y665F variant of mouse *Stat5b* protects against acute kidney injury through transcriptomic shifts in renal gene expression"

Supplementary Figure 1

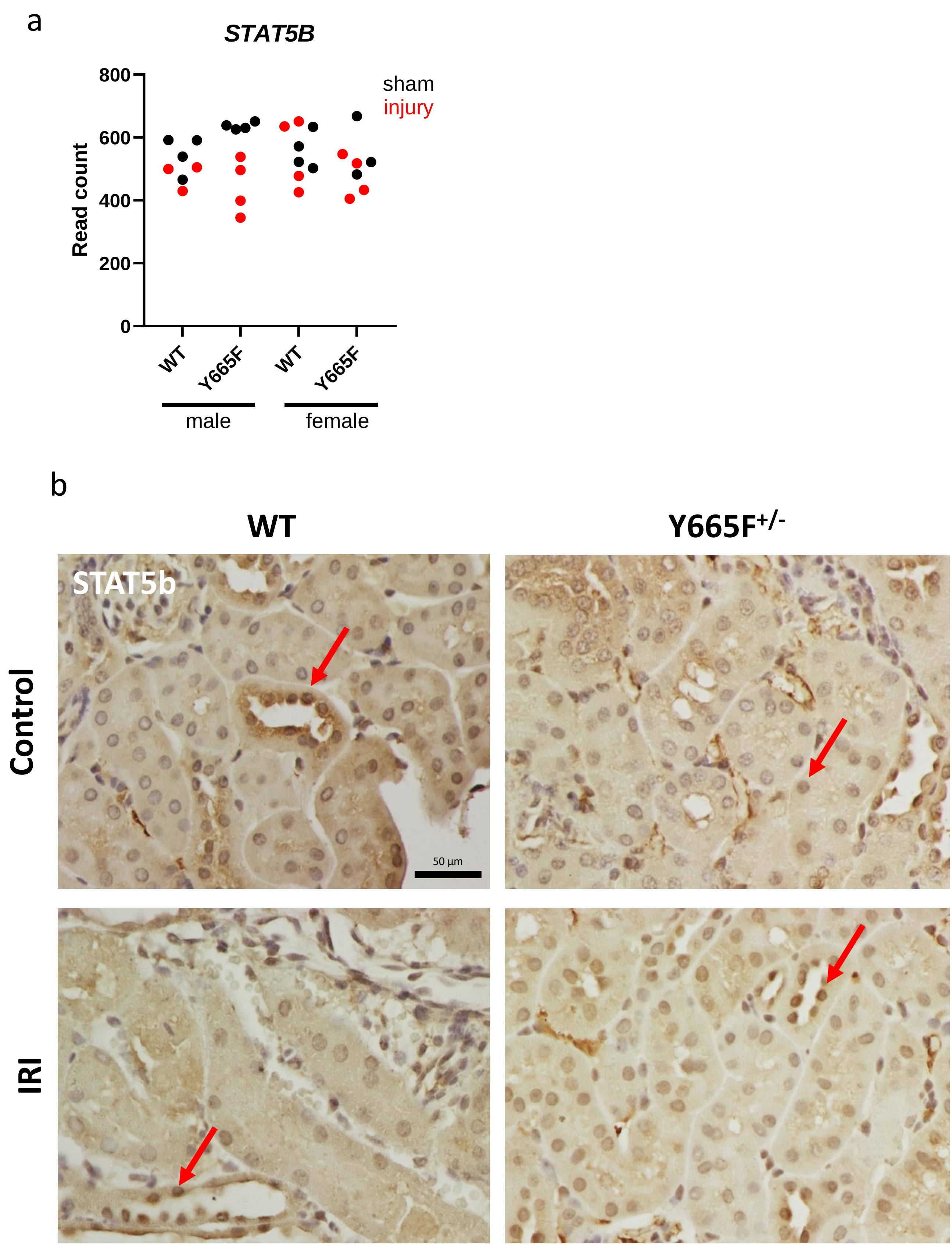


**Supplementary Fig. 1** Normalized DeSeq2 reads of *Stat5b* in male and female wild type and Y665F mutants before and after injury (a), n = 3-4. Representative STAT5b staining images of renal tissue from male wild type and Y665F mice at the baseline and 24 hours after injury (b); bar = 50 µm, 400x magnification, arrows point to examples of positive nuclei.

Supplementary Figure 2

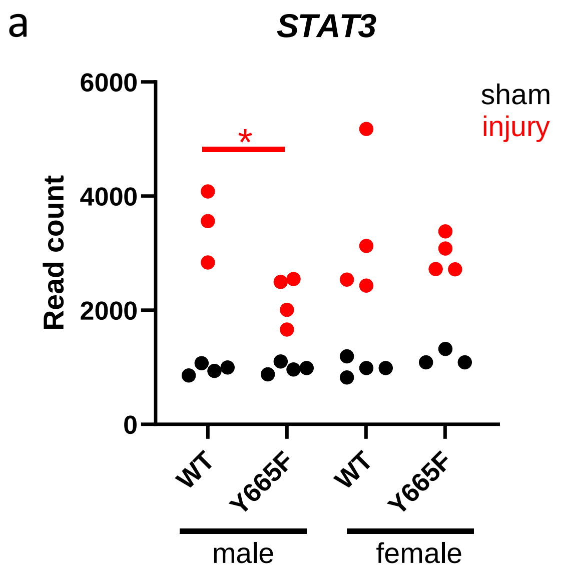


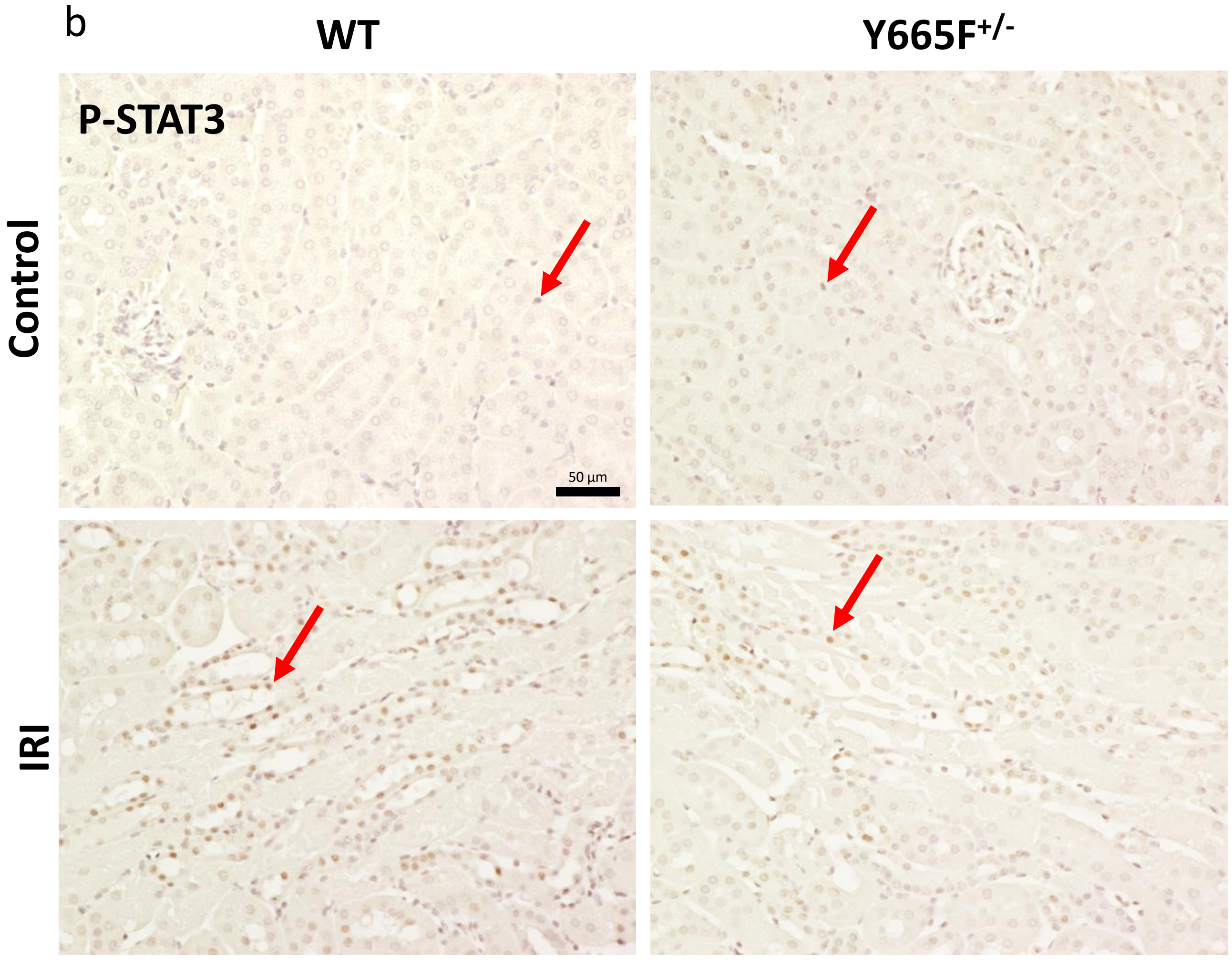


**Supplementary Fig. 2** Normalized DeSeq2 reads of *Stat3* in male and female wild type and Y665F mutants before and after injury (a), n = 3-4; * $P < 0.05$. Representative P-STAT3 staining images of renal tissue from male wild type and Y665F mice at the baseline and 24 hours after injury (b); bar = 50 µm, 400x magnification, arrows point to examples of positive nuclei.

Supplementary Figure 3

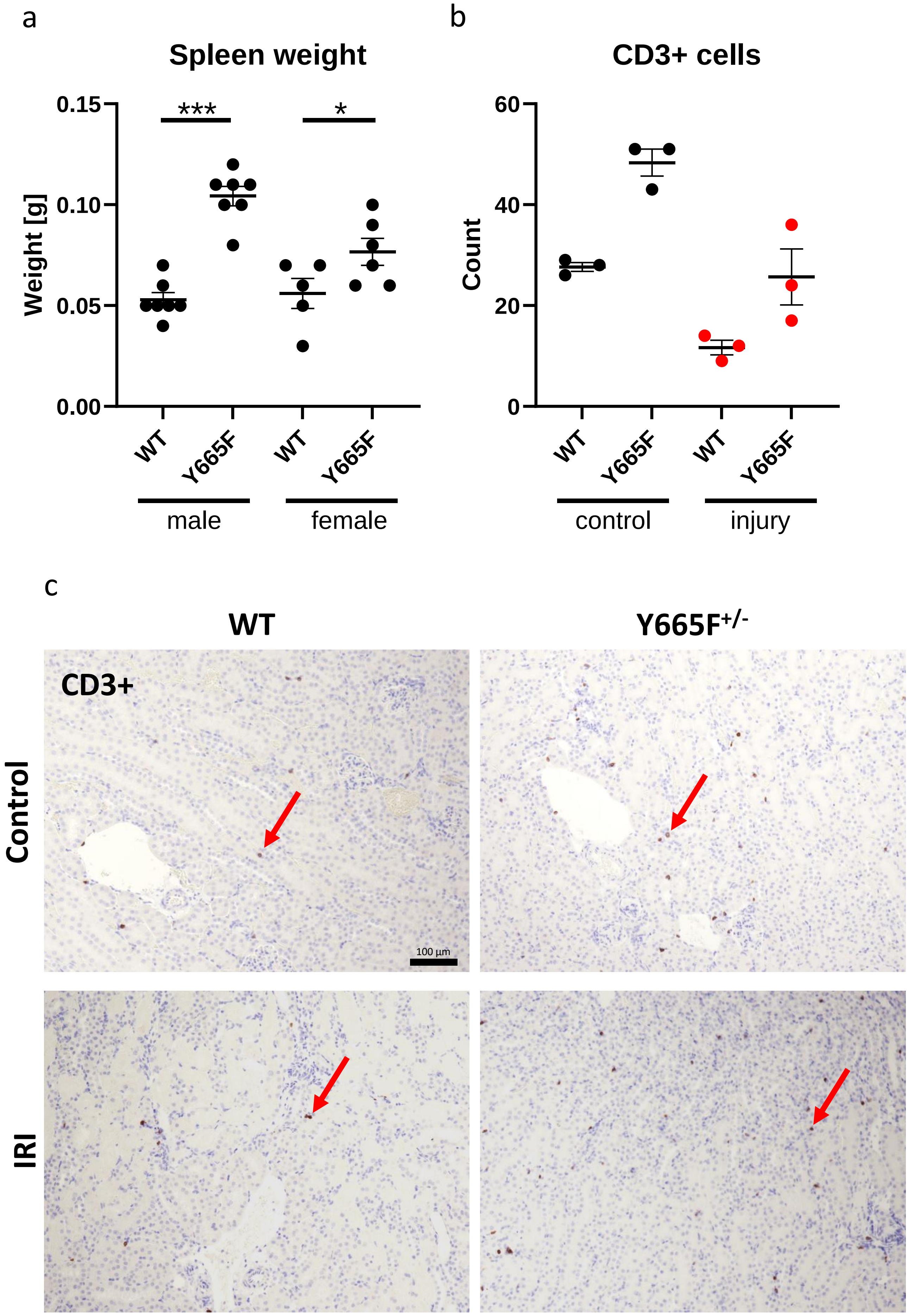


**Supplementary Fig. 3** Spleen wights of male and female wild type and Y665F mice (a). n = 5-7, Bar = SEM, Two-way ANOVA with group mean comparisons, ** *P* < 0.01, *** *P* < 0.01. Number of CD3-positive cells per random microscope fields (b), n=3. Representative CD3 staining images of renal tissue from male wild type and Y665F mice at the baseline and 24 hours after injury (b); bar = 100 µm, 200x magnification, arrows point to examples of positive cells.

Supplementary Figure 4

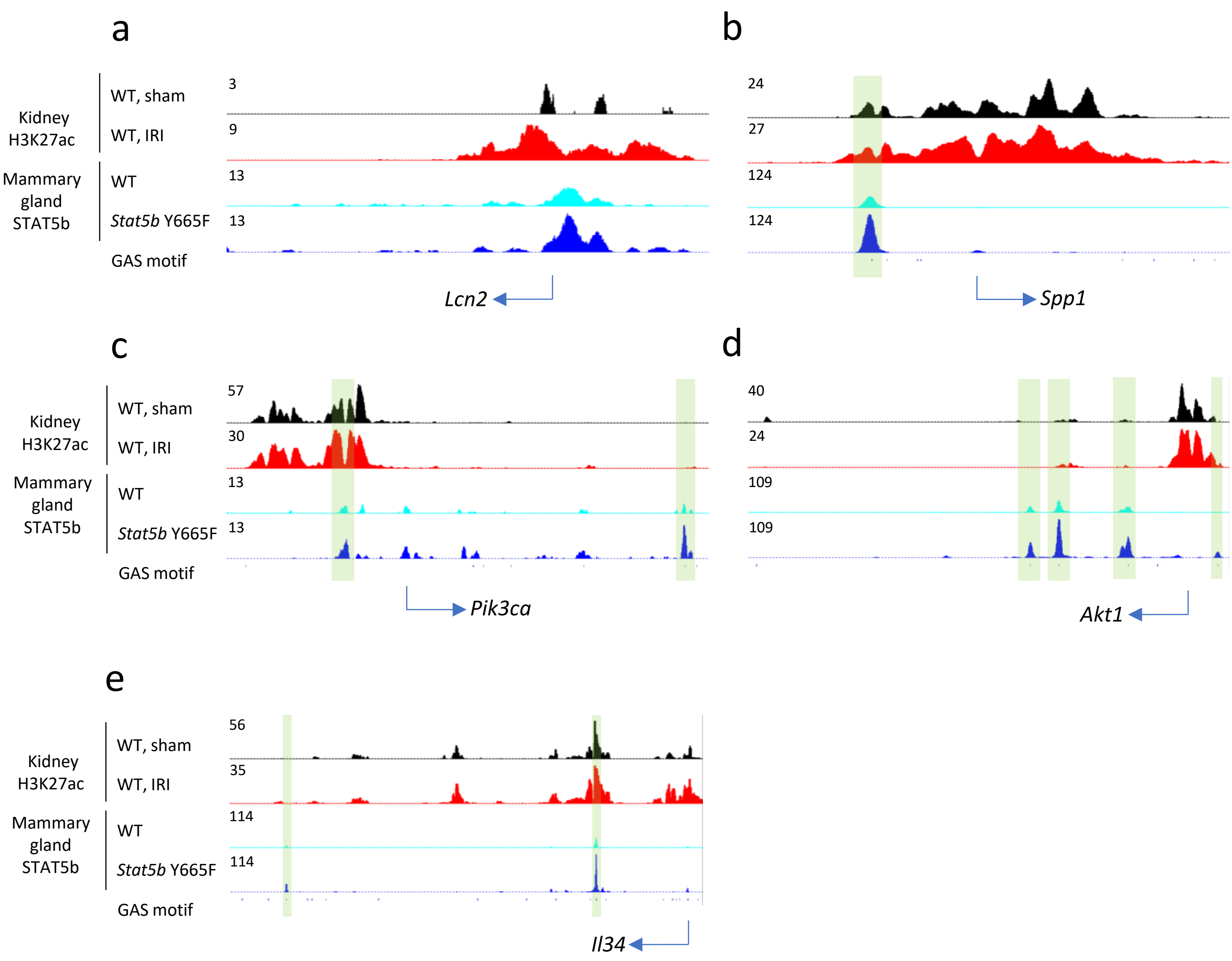


**Supplementary Fig. 4** ChIP-seq tracks visualizing H3K27ac (active chromatin) in renal tissue in wild type sham and AKI mice, as well as mammary gland STAT5b binding in late pregnancy (P18) in wild type and homozygous *Stat5b*$^{Y665F}$ mice. Relevant GAS (STAT-binding, TTCnnnGAA) motifs and potential STAT5b binding sites are marked with semi-transparent shading.

Supplementary Figure 5

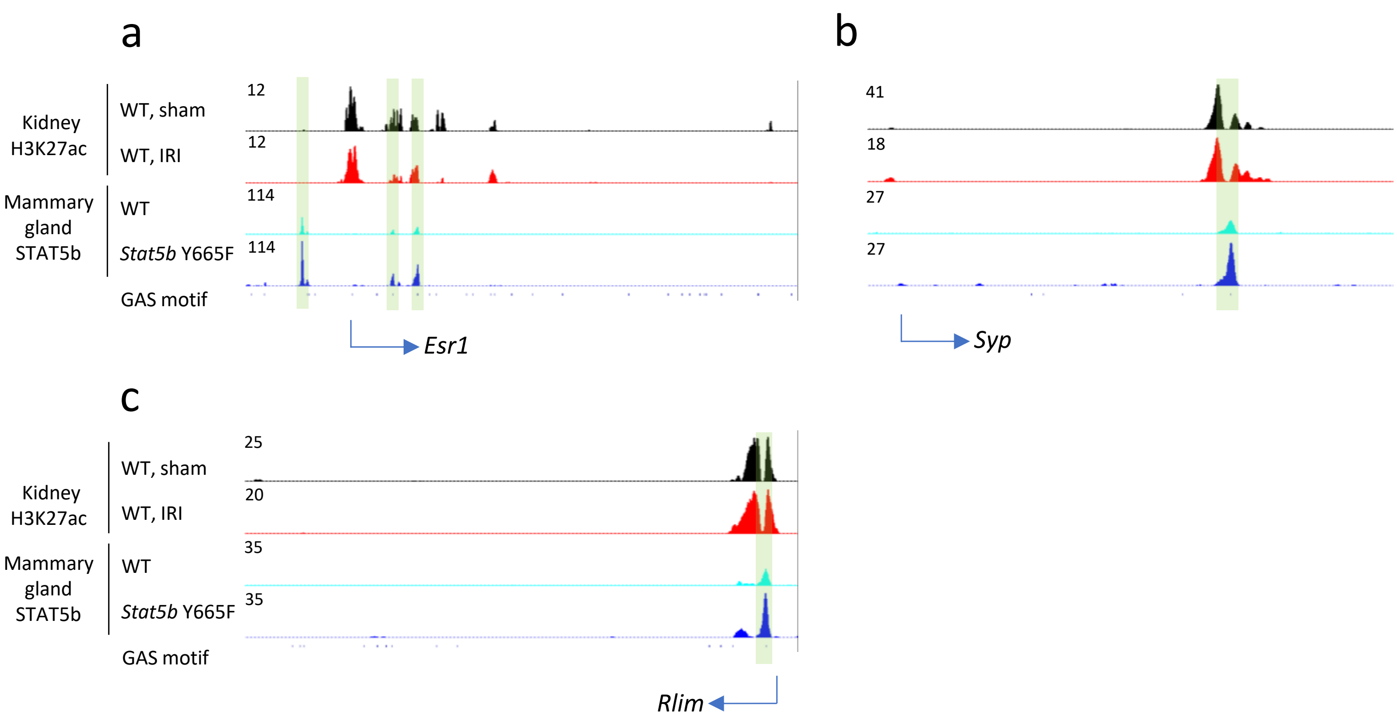


**Supplementary Fig. 5** ChIP-seq tracks visualizing H3K27ac (active chromatin) in renal tissue in wild type sham and AKI mice, as well as mammary gland STAT5b binding in late pregnancy (P18) in wild type and homozygous *Stat5b*$^{Y665F}$ mice. Relevant GAS (STAT-binding, TTCnnnGAA) motifs and potential STAT5b binding sites are marked with semi-transparent shading.
